# Exploration of the intricacies of low light-induced changes in cigar tobacco leaf anticlinal growth: A holistic approach from anatomical and hormonal levels to gene expression

**DOI:** 10.1101/2023.12.20.572520

**Authors:** Xinghua Ma, Jinpeng Yang, Xiaochun Ren, Keling Chen, Chunlei Yang, Huajun Gao, Rayyan Khan

## Abstract

Cigar tobacco stands as a pivotal economic crop, with its leaf growth and development profoundly influenced by light intensity. It specifically aims to investigate how leaf morphology and anticlinal growth respond to varying light intensities, including normal light intensity (NL–300 μmol m^−2^ s^−1^) and lower light intensity (LL–100 μmol m^−2^ s^−1^). The research elucidates significant morphological shifts in cigar tobacco leaves under LL, revealing notable alterations in leaf area, leaf length, and leaf width. Early reductions in leaf dimensions, ranging from 30% to 48%, were succeeded by a substantial enhancement in expansion rates from day 9 to day 26, contributing to expanded leaf surfaces at later stages. Upper epidermis thickness declined by 29-19%, with a notably slower expansion rate in the initial 20 days. Palisade cell length consistently decreased by 52-17%, corresponding with upper epidermis trends. Spongy tissue thickness was reduced by 31-12%, with a slower expansion rate in LL for the initial 14 days, and leaf thickness dropped by 34-11%. LL resulted in slower leaf anticlinal expansion, leading to reduced leaf thickness (LT). LL significantly influenced phytohormones in cigar tobacco leaves. Gibberellic acid (41% to 16%) and auxin (20% to 35%) levels were found in higher amounts, while cytokinin levels (19% to 5%) were lowered compared to NL, indicating the intricate regulatory role of light in hormonal dynamics. The observed increase in LT and different cell layers at specific time points (day 8, day 12, day 24, and day 28) under LL, although lower than NL, may be attributed to elevated expression of genes related to cell expansion, including *GRF1*, *XTH*, and *SAUR19* at those time points. This comprehensive understanding elucidates the intricate mechanisms by which light intensity orchestrates the multifaceted processes governing leaf anatomy and anticlinal expansion in cigar tobacco plants.

## Introduction

Light intensity emerges as a pivotal environmental factor profoundly influencing plant growth and morphogenesis (Sassi et al. 2013). When subjected to low light levels, plants exhibit a notable response by reducing the thickness of their leaves (Wei et al. 2023). Research highlights that low light intensity correlates with an increase in leaf area (Tang et al. 2022). Additionally, low light intensity also hinders the photosynthetic capacity of plant leaves and reduces their capacity to accumulate photosynthetic materials (Xiang et al. 2021). Furthermore, the effects of low light intensity on plant growth encompass various aspects, including the elongation of hypocotyls, internodes, and petioles, a reduction in branching, and alterations in leaf mass. These changes are integral components of the shade avoidance response triggered by low light intensity (Nozue et al. 2015; Tang et al. 2021). Therefore, gaining a comprehensive understanding of the intricate interplay between light intensity and leaves holds paramount importance for enhancing crop yield and quality, especially for crops harvested for their leaves.

Plant leaves, originating from pluripotent stem cells within the stem apical meristem, manifest as differentiated organs regulated by a sophisticated genetic network. This network intricately coordinates cell proliferation and differentiation, guiding organized growth along the transverse, longitudinal, and anticlinal axes that define leaf structure (Nikolov et al. 2019; Satterlee and Scanlon 2019). Leaf thickness (LT), indicating the distance from upper to lower surfaces (Coneva and Chitwood 2018), determines the optical route of light absorption, reflection, or transmission. LT exhibits correlations with various environmental variables. Previous studies, including those by (Li et al. 2014) and (Murchie et al. 2005), have indicated an increase in LT in response to high-light conditions. The regulation of growth processes at the tissue level emerges as a crucial aspect of leaf growth regulation. John et al. (2013) suggested the thickness of mesophyll tissue as a pivotal component of overall LT. The most significant factor leading to an increase in LT is the expansion of palisade mesophyll cells in the dorsoventral direction (Coneva et al. 2017). Remarkably, low light intensity detrimentally affects mesophyll tissue thickness, particularly in palisade and spongy tissues (Zhang et al. 2022).

The intricate processes of plant growth and development are closely intertwined with various factors, with phytohormones playing a central role and carrying out specific functions. Essential plant hormones, such as auxin, cytokinin, and gibberellins, actively participate in steering the regulation of leaf development, as highlighted in the works of researchers like Robil and McSteen (2023), Hussain et al. (2021), and Mu et al. (2018). These scholars have provided insights into the complex interplay of these hormones, particularly in the processes of leaf formation, expansion, and size determination during leaf morphogenesis. The intricate orchestration of plant growth and development in response to varying light conditions hinges on fluctuations in hormone levels and corresponding signals (De Wit et al. 2016). Beyond hormones, the role of gene expression is also integral to a plant’s adaptation to light-intensity variations (Wu et al. 2017). It’s important to recognize that leaf development is the outcome of a complex interplay between genes and hormones (Du et al. 2018). Consequently, unraveling the disparities in hormone levels and gene expression under low light intensity serves as a foundational step in comprehending the growth and development of plant leaves.

Cigar, an important cash crop, is cultivated extensively worldwide. A cigar consists of three essential components: the wrapper, binder, and filler tobacco. Of the three components, the wrapper tobacco constitutes the smallest proportion, yet commands the highest unit price. The birthplace of cigars is Cuba, but presently, cigar tobacco raw materials are primarily concentrated in Central America, the Caribbean, central and western Africa, and Southeast Asia. The unique requirements for growing conditions have limited the widespread cultivation of cigar wrapper tobacco. Given that superior cigar wrappers necessitate leaves of moderate size, thin thickness, and delicate texture, they are typically cultivated under shade conditions (Wu et al. 2021). Consequently, investigating the growth and development disparities of cigar wrapper tobacco under varying light intensities represents a crucial step in enhancing cigar wrapper quality.

While previous studies have elucidated disparities in leaf development, hormone levels, and gene expression in certain plant species under varying light intensities, these investigations predominantly focused on specific timer frames and lack to comprehensively examine the entire leaf development process. Currently, there remains a lack of clarity regarding how cigar wrapper development differs under low light conditions and when these differences manifest throughout the developmental timeline under varying light intensities. To bridge this knowledge gap, our approach involves continuous sampling to achieve two primary objectives: Firstly, we aim to scrutinize the evolving dimensions of cigar leaves, encompassing changes in length, width, and thickness. Secondly, we seek to explore variations in hormone concentrations and gene expression levels. The outcomes of this study are expected to enhance our comprehension of how cigar leaf development responds to light intensity, providing a valuable theoretical foundation for the cultivation of cigar wrapper tobacco.

## Materials and methods

### Plant material and treatments

The cigar tobacco *Nicotiana tabacum* L. cv. Guyin 4 was generously provided by the Haikou Cigar Research Institute of the China National Tobacco Corporation (Hainan, China). A potted experiment was conducted within an artificial climate chamber Jimo Experimental Base of the Tobacco Research Institute of the Chinese Academy of Agricultural Sciences (Qingdao, China). The cultivation of Guyin 4 began with the tray cultivation method under the light intensity of 300 ± 12 μmol m^−2^ s^−1^. Once the tobacco seedlings had reached the stage of 6 leaves, we carefully selected robust and uniform tobacco plants. The carefully selected plants were transplanted into plastic pots with an 18 cm diameter. The pots were filled with a substrate composed of peat soil, vermiculite, and perlite in a 3:1:1 ratio (V/V/V). During the experiment, two distinct light intensities were used, namely, normal light intensity (NL—300 μmol m^−2^ s^−1^) and low light intensity (LL—100 μmol m^−2^ s^−1^). This experiment was carried out within a controlled environment that featured a 14-hour day period at 28 ℃ and a 10-hour night period at 23 ℃ while maintaining a relative humidity level of 70±5%. To control the light intensity, we utilized LED red and blue light sources. Our measurements and sampling commenced when the tenth leaf emerged from the apical bud.

### Leaf growth analysis

On a daily basis, we used a ruler to measure both leaf length and leaf width (n=3 plants). Subsequently, we calculated the leaf area for the measured leaves using the formula: Leaf area = leaf length × leaf width × 0.6345 (Liu 1996). To evaluate the proportional expansion rates, we converted the average measurement values of leaf length, width, and area into logarithmic values. Subsequently, these transformed values underwent local fitting through a 5-point quadratic function. From this fitted function, we derived the first derivative, which allowed us to calculate the relative expansion rates (Kalve et al. 2014).

### Anatomical study

The leaf samples were collected at various time points: 2 days (d), 4d, 6d, 8d, 10d, 12d, 14d, 16d, 20d, 24d, and 28d (n=3 leaves). The sampling method involved puncturing the leaf blade at the center with a 1.25 cm diameter while avoiding veins. Subsequently, these samples were preserved in an FAA solution, which consisted of 38% formaldehyde, glacial acetic acid, and 70% alcohol in a 5:5:90, v/v/v ratio. The fixed leaf fragments underwent dehydration using a gradient ethanol solution series (75%– 100%) and subsequently rinsed in xylene. Following dehydration, the leaf samples sections were embedded in paraffin. Afterward, the samples were stained with a Safranin O solution (1%) and Fast Green FCF (0.5%). The observations were carried out using the Leica DM 2000 microscope (Leica, Wetzlar, Germany) coupled with a Leica DMC 2900 camera (Leica, Wetzlar, Germany), allowing us to capture microscopic images for analysis. To evaluate the samples comprehensively, we quantified the number of cell layers and precisely measured various aspects, including the overall leaf thickness, as well as the thickness of the upper and lower epidermis, palisade tissue, and spongy tissue. These measurements were made using Image J software (http://rsbweb.nih.gov/ij/). Moreover, the mean values for LT and the thickness of individual cell layers underwent a logarithmic transformation. These transformed values were locally fitted using a 5-point quadratic equation function, and their respective expansion rates were computed using the first derivative (Kalve et al. 2014).

### Phytohormone determination

Leaf samples were collected at distinct time intervals: 4d, 8d, 12d, 16d, 20d, 24d, and 28d. Following their collection, the leaf samples were rinsed in liquid nitrogen and subsequently stored at -80°C. The contents of gibberellic acid, auxin, abscisic acid, and cytokinin (n=3 biological repeats) were determined following the guidelines of ELISA kits (Cat No. 2Pl-KMLJ91022p, Cat No.2Pl-KMLJ91127p, Cat No.2Pl-KMLJ91056p, and Cat No. 2Pl-KMLJ91106p), respectively, purchased from Nanjing Camilo biological engineering Co., Ltd. (www.njkmlbio.com) (Nanjing, China).

### RNA extraction and gene expression analysis

Total RNA was extracted from the frozen leaf samples (n=3 biological repeats) following the guidelines of the FastPure Plant Total RNA Isolation Kit (Vazyme Biotech Co., Ltd, Nanjing, China) which were collected from various light intensity treatments at 4d, 8d, 12d, 16d, 20d, 24d, and 28d time points. Subsequently, we evaluated both the concentration and quality of the RNA. This assessment was carried out using a Nanophotometer (Implen) for concentration measurement and agarose gel electrophoresis for quality analysis. Next, we synthesized the first-strand cDNA using the PrimeScript 1st-strand cDNA synthesis kit (TaKaRa, Japan). The qRT-PCR was conducted in a 20μl reaction volume employing SYBER Green Master Mix (TaKaRa, Japan). The qRT-PCR was carried out using the Applied Biosystems QuantStudio3 real-time PCR machine (Thermo Fisher, Waltham, MA, USA). For data analysis, the *Actin* gene served as an internal reference gene, and the data were analyzed using the 2^−ΔΔCT^ method (Livak and Schmittgen 2001). All the primer pairs used for gene expression analysis are presented in Table 1.

**Table 1.**
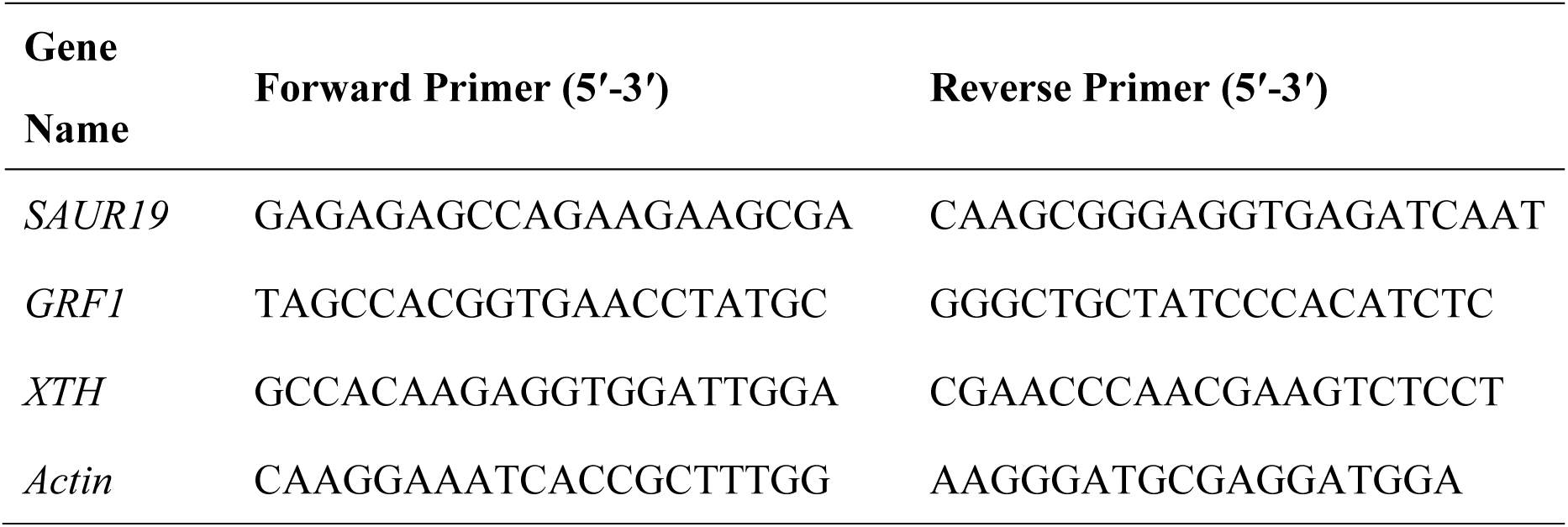
Primers used for RT-qPCR.

### Data analysis

We used Statistix 8.1 software (Analytical Software, Tallahassee, FL, USA) to analyze the collected data with a one-way ANOVA model. An LSD test was used to determine significant differences between the mean values of NL and LL treatments, with a significance level set at p < 0.05 and p < 0.01 Subsequently, we presented the data graphically using Origin 2018 software (OriginLab Corporation, Northampton, MA, USA) and GraphPad Prism 8.0 (GraphPad Software, San Diego, California USA) software.

## Results

### Changes in leaf morphology and expansion rates

The effect of low light intensity on leaf morphology revealed statistically significant effects (p < 0.05 and p < 0.01) on leaf area (LA), leaf length (LL), and leaf width (LW) over the 28-day observation period (Fig. 1). In the initial 11 days, LL showed fold changes ranging from 0.67 to 0.52, indicating a reduction between 33% and 48% compared to normal light (Fig. 1A). From day 12 to day 20, LL fold changes ranged from 0.71 to 0.97, representing a decrease of 3% to 29% under low light. Subsequently, from day 23 to day 28, LL exhibited an increase in fold changes (1.05 to 1.10), indicating a 5% to 10% enhancement under low light (Fig. 1A). The LL expansion rate under low light surpassed that under normal light from day 11 to day 26 (Fig. 1B). LW displayed a comparable trend to LL under low light, with fold changes ranging from 0.56 to 0.70 (a decrease of < 44% and > 30%) during the initial 10 days compared with normal light (Fig. 1C). From day 11 to day 18, LW fold changes ranged from 0.74 to 0.97 (a reduction of 26% and 3%) under low light compared with normal light. Additionally, an increase in LW values (fold change of 1.04—4% increase to fold change of 1.07—7% increase) was observed between day 22 and day 28 under low light (Figure 1C). The LW expansion rate was higher in low-light plants than in normal ones between day 9 and day 26 (Fig. 1D), resulting in broader leaves in the later days (Fig. 2). Furthermore, LA showing fold changes < 0.60 and 0.40 (a reduction between 40% and 60%) in the initial 12 days under low light (Fig. 1E). Between day 13 and day 18, LA values under low light exhibited a fold change of 0.64 (reflecting a 36% reduction) to 0.90 (indicating a 10% reduction). In contrast, there was an increase in fold changes ranging from 1.05 (5% increment) to 1.17 (17% increment) in comparison to normal light conditions from day 21 to day 28 (Fig. 1E). This ultimately led to a higher leaf surface expansion rate in low light plants compared to normal ones from day 9 to day 26 (Fig. 1F), contributing to the development of larger leaves (Fig. 2).

**Fig. 1.**
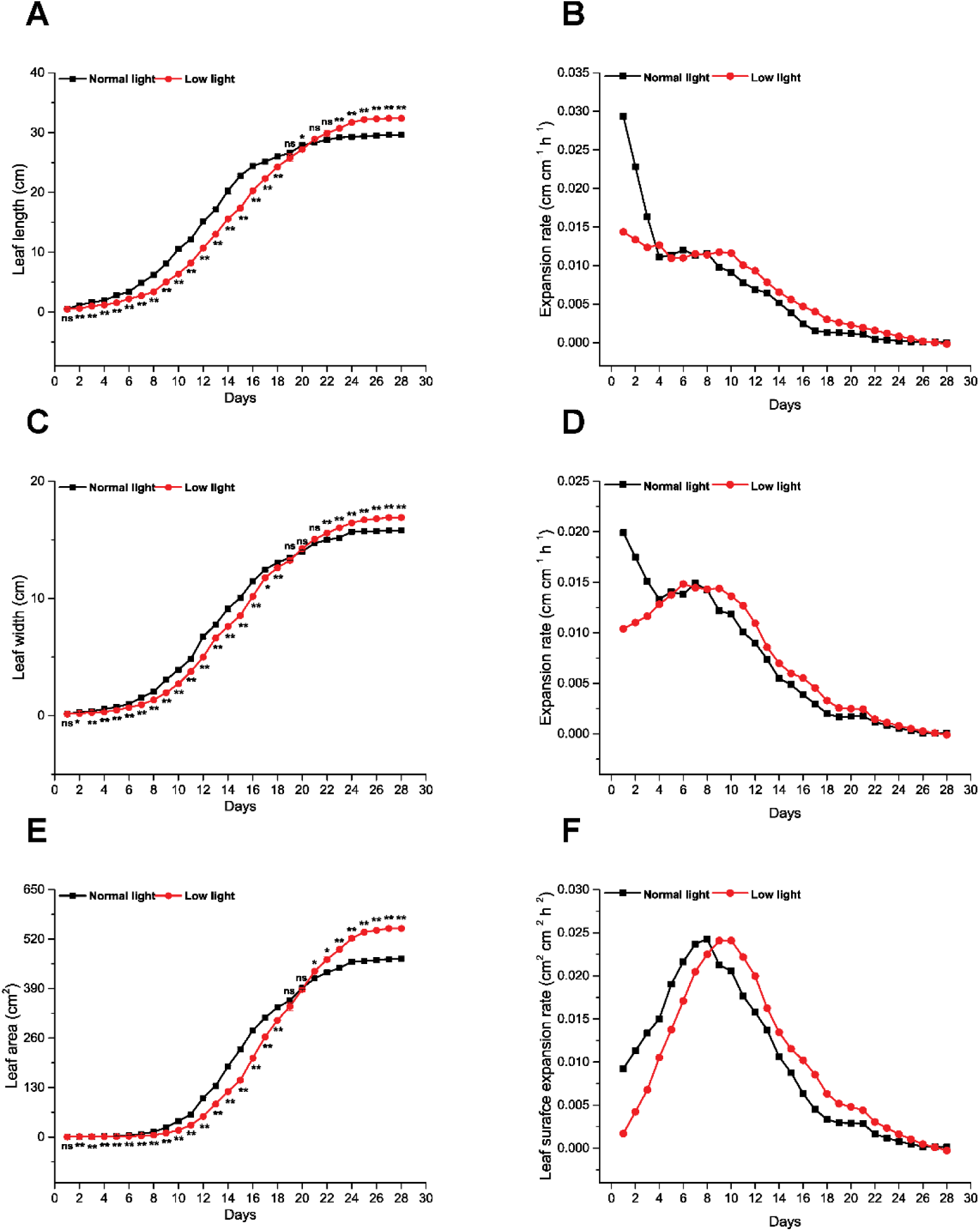
Effect of low light intensity on cigar tobacco leaf morphology. This figure outlines changes in leaf length (**A**), leaf length expansion rate (**B**), leaf width (**C**), leaf width expansion rate (**D**), leaf area (**E**), and leaf surface expansion rate (**F**) under both low light and normal light conditions over the 28-day observation period. The data is presented as mean ± SE at each time point, with * and ** indicating significant differences at p < 0.05 and p < 0.01, respectively, while ns represents non-significant differences.

**Fig. 2.**
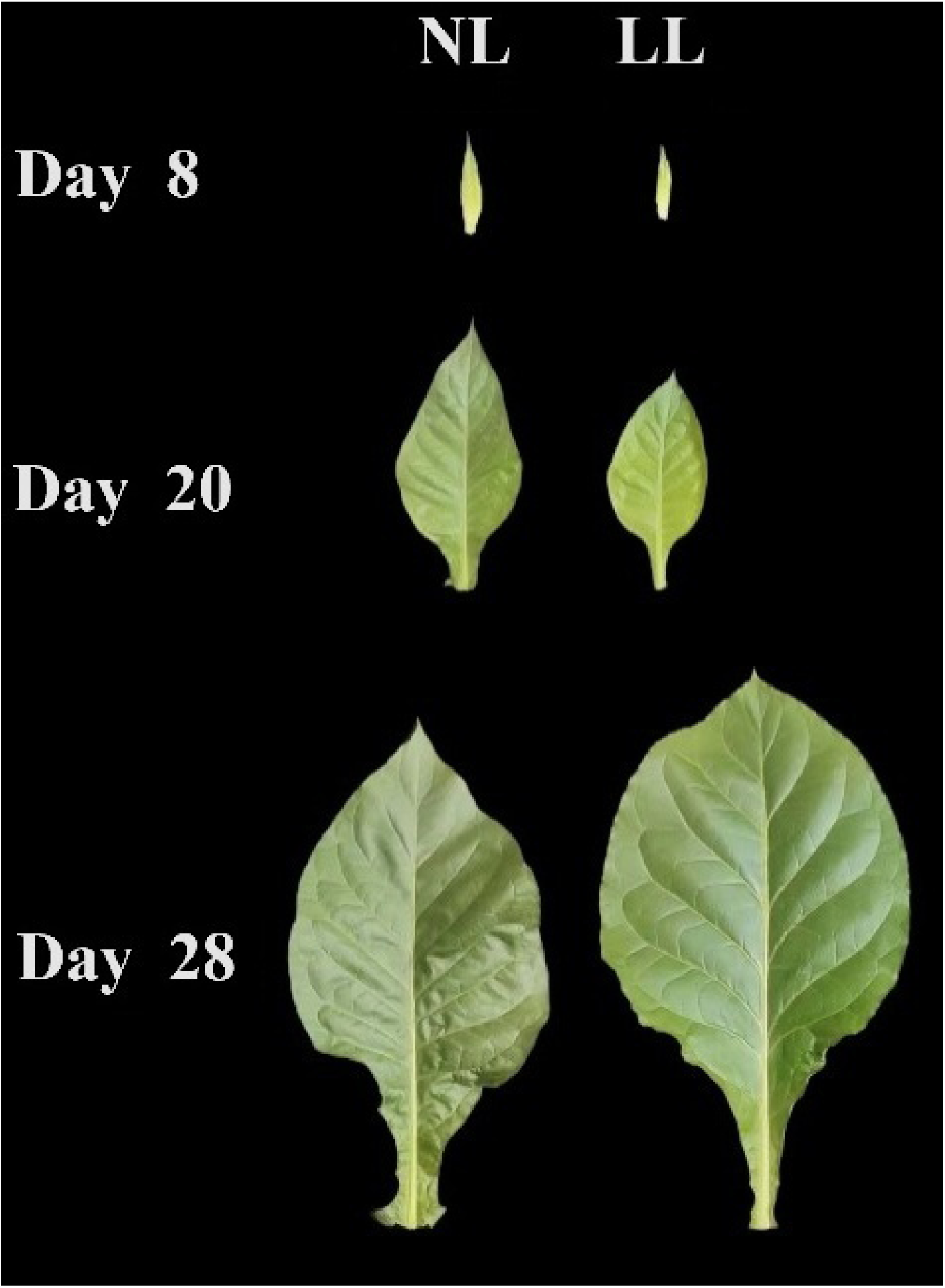
Representative images of cigar tobacco leaf as affected by low light and normal light at day 8, day 20, and day 28. NL represents normal light intensity (300 μmol m^−2^ s^−1^) and LL represents low light intensity (100 μmol m^−2^ s^−1^).

### Changes in the leaf anatomy and anticlinal expansion rate

The cross-sectional analysis of cigar tobacco leaves revealed an absence of differentiation among distinct cell layers (upper epidermis—UE, palisade cells length—PCL, spongy tissues thickness—STT, and lower epidermis—LE) up to 8 days under normal light intensity (NL) and 10 days under low light intensity (LL) (Fig. 3). Beyond these time points, clear distinctions emerged in the cell layers, featuring upper and lower epidermis as well as mesophyll cell layers including palisade and spongy cells (Fig. 3). From the beginning, there were seven cell layers in normal light and six cell layers in low light plants. Upon day 8 and day 10, there was a clear differentiation between the cell layers which could be differentiated into UE, PC, SC, and LE under NL and LL, respectively, but there was a difference in the spongy cell layers in NL (4 cell layers) and LL (3 cell layers) (Fig. 3). Notably, there was a discernible reduction in leaf thickness (LT) influenced by low light.

**Fig. 3.**
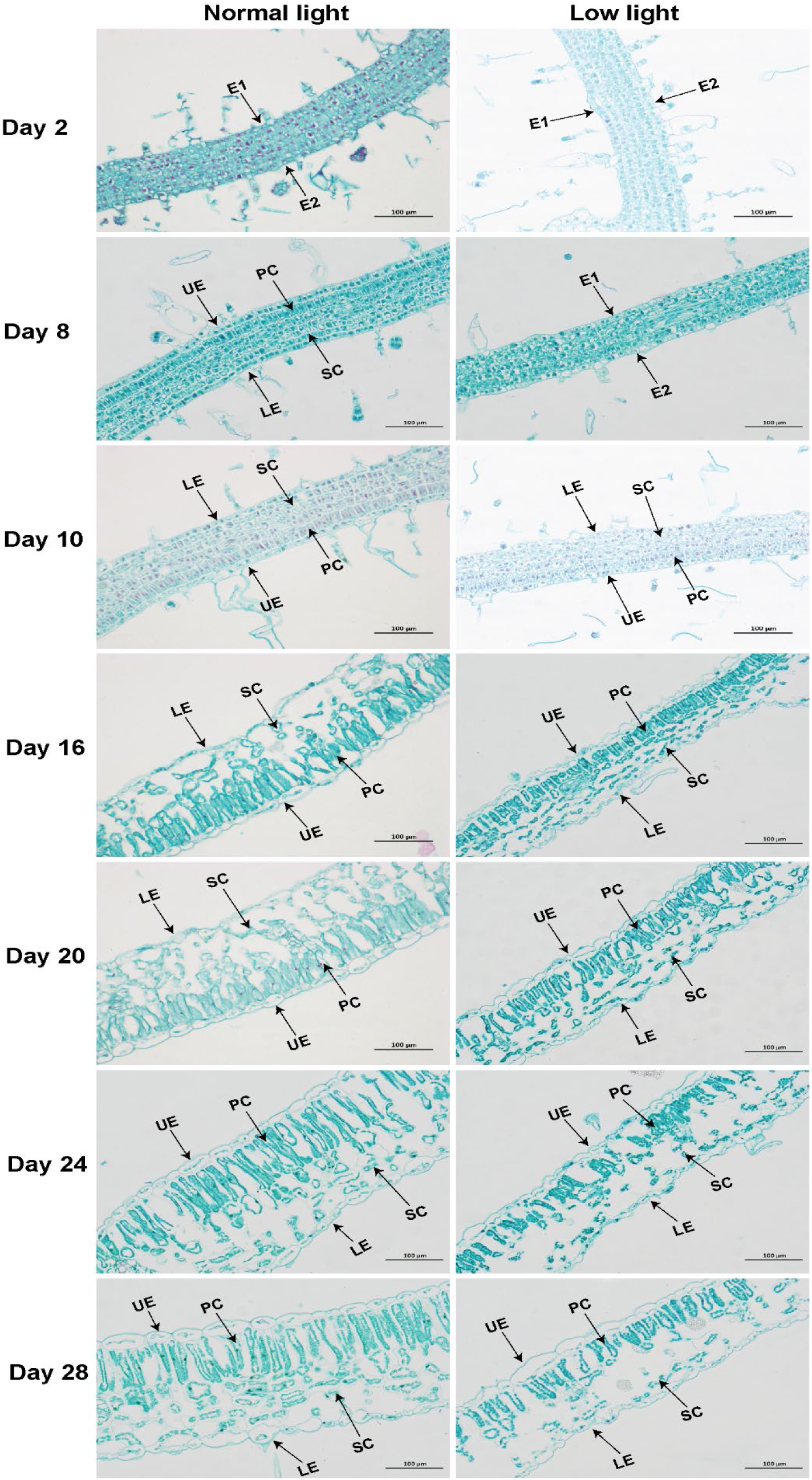
Effect of low light on leaf structure. Representative images depicting leaf structure under normal light and low light conditions at various time points. E1— epidermal cell layer 1, E2—epidermal cell layer 2, UE—upper epidermis, PC— palisade cells, SC—spongy cells, and LE—lower epidermis.

Low light significantly (p < 0.01) affected UE, PCL, STT, LE, and LT. The upper epidermis thickness was noticeably thinner in LL, exhibiting a fold change between 0.71 and 0.81, representing a reduction between 29% and 19% over the 28-day observation period compared with NL (Fig. 4A). The expansion rate of the upper epidermis was slower under LL, ranging from triple to septuple times, up to 20 days, and surpassed thereafter (Fig. 4B). PCL was consistently shorter under LL throughout the observation period, with a fold change ranging from 0.48 to 0.83 (a reduction of 52% to 17%) compared with NL (Fig. 4C). The PCL expansion rate showed a similar trend to UE up to 20 days and was one to triple times slower in LL compared to NL (Fig. 4D). Furthermore, STT was lower in LL compared to NL, with a fold change spanning from 0.69 (31% reduction) to 0.88 (12% reduction) throughout the entire observation period (Fig. 4E). The STT expansion rate was double times slower in LL at day 10, day 12, and day 14, equal at day 16 and day 20, and subsequently faster compared with NL (Fig. 4F). Low light leaves exhibited a thin LE, with a fold change between 0.84 (16% reduction) and 0.87 (13% reduction) compared with NL (Fig. 4G). The LE expansion rate was one to double times slower in LL up to 16 days and almost equal at day 20, day 24, and day 28 compared with NL (Fig. 4H). Finally, LT showed a reduction of < 6% (fold change from 0.94 to 0.96) in the first 10 days. However, afterward, LT was thinner, displaying a fold change ranging from 0.66 (34% reduction) to 0.89 (11% reduction) under LL compared with NL (Fig. 4I). The leaf anticlinal expansion rate in LL was slower, ranging from one to quadruple times, and almost equal afterward compared with NL (Fig. 4J). The leaf anticlinal expansion rate and the expansion rate of each tissue layer were observed to be lower under LL for the initial 20 days compared to NL, resulting in a reduction in leaf thickness by the end of the observation period.

**Fig. 4.**
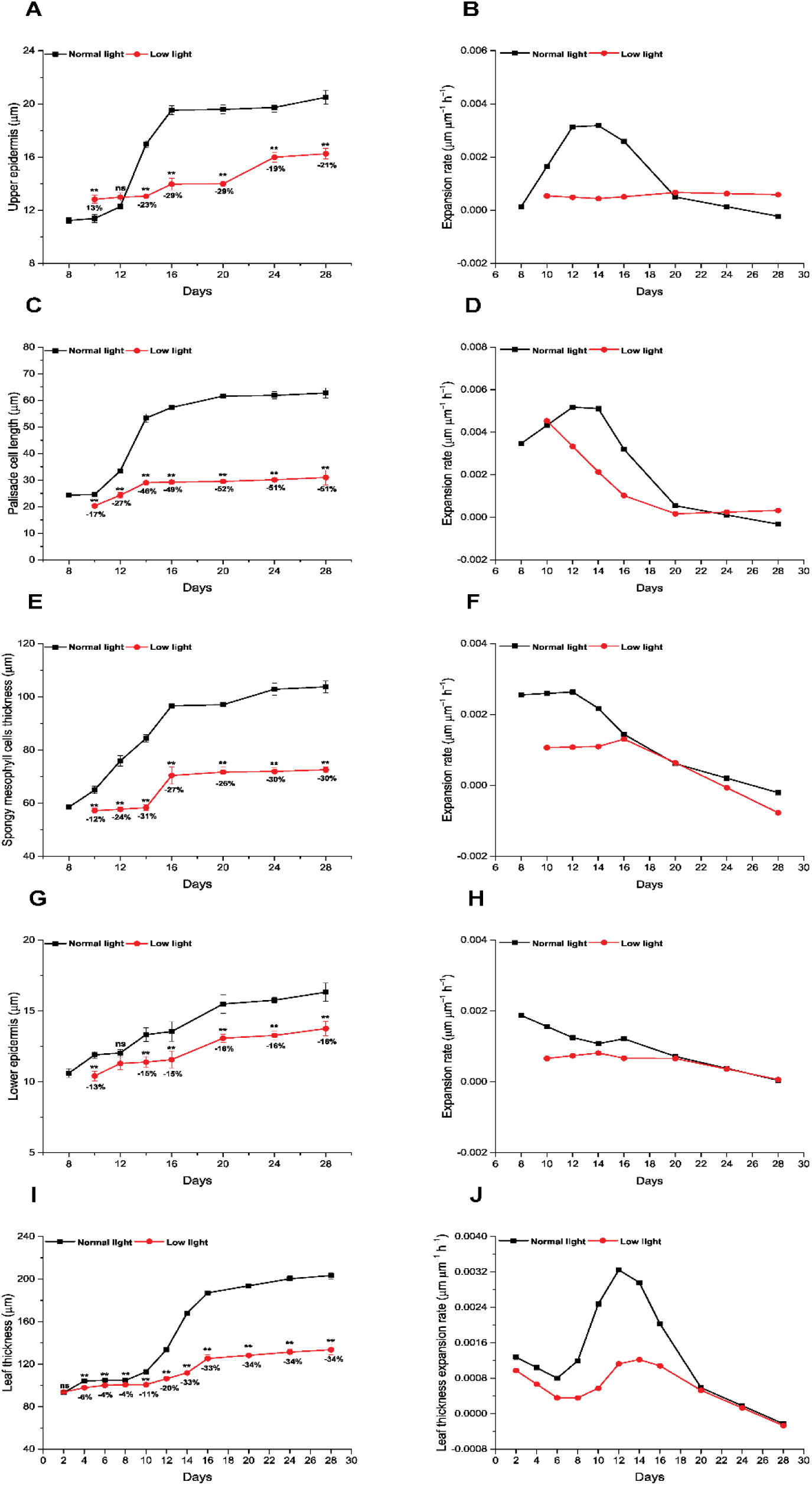
Impact of reduced light on leaf thickness, individual tissue layer thickness, and their corresponding expansion rates. Parameters include upper epidermis thickness (**A**), upper epidermis expansion rate (**B**), palisade cell length (**C**), palisade cell expansion rate (**D**), spongy tissue thickness (**E**), spongy tissue expansion rate (**F**), lower epidermis thickness (**G**), lower epidermis expansion rate (**H**), leaf thickness (**I**), and leaf anticlinal expansion rate (**J**) under low light and normal light conditions over the 28-day observation period. Data are presented as mean ± SE at each time point, with ** indicating significant differences at p < 0.01, and ns representing non-significant differences.

### Changes in phytohormone content

Low light showed a significant (p < 0.05 and p < 0.01) effect on the content of phytohormones, including gibberellic acid (GA), indole acetic acid (IAA), abscisic acid (ABA), and cytokinin (CK) (Fig. 5). GA content consistently exhibited higher levels under LL across the 28-day observation period, reaching its peak with a 41% increase at day 4 and a minimum of 16% at day 28 compared to NL (Fig. 5A). IAA content showed elevation under LL, especially in the early stages, with increments of 35% each at day 4 and day 8, persisting until day 20, while no significant changes were observed at day 24 and day 28 compared to NL (Fig. 5B). Interestingly, ABA content increased only in the final days of leaf growth, registering a 5% and 8% rise at day 20 and day 28 compared to NL (Fig. 5C). Conversely, CK content exhibited an opposing trend throughout the growth period, declining up to day 20 and experiencing a more substantial reduction of 19% and 18% at day 4 and day 8, followed by a milder reduction of 5% and 9% at day 20 and day 24 compared to NL (Fig. 5D).

**Fig. 5.**
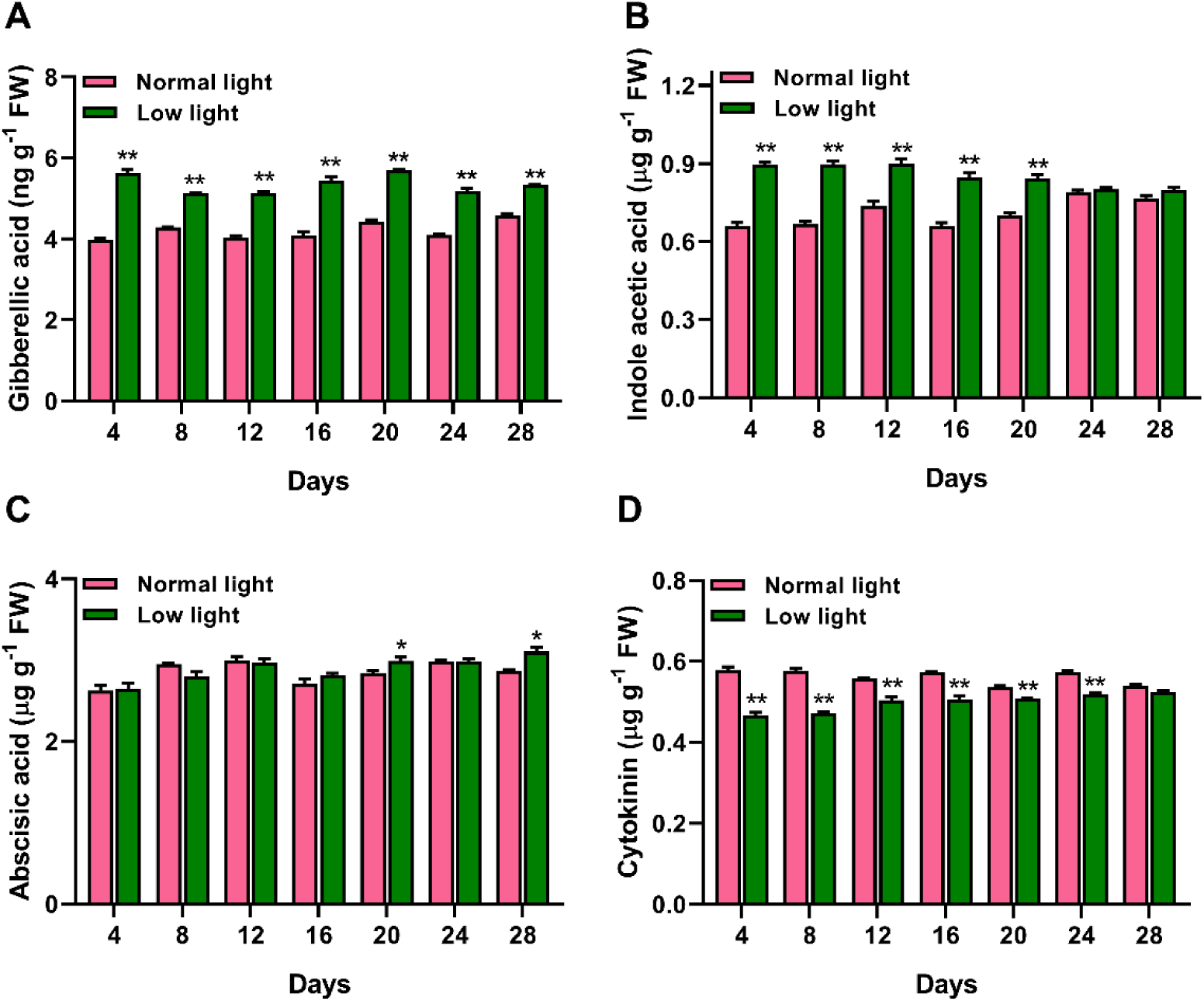
Influence of low light on phytohormones content. Gibberellic acid content (**A**), Indole acetic acid content (**B**), Abscisic acid content (**C**), and Cytokinin content (**D**) as affected by low light and normal light over the 28-day observation period. Data are presented as mean ± SE at each time point, with * and ** indicating significant differences at p < 0.05 and p < 0.01, respectively, while ns representing non-significant differences.

### Changes in gene expression levels related to cell proliferation and expansion

Low light significantly influenced gene expression levels related to cell expansion, including *GRF1*, *XTH*, and *SAUR19* (Fig. 6). *GRF1* exhibited upregulation in LL at day 8 (3.5-fold change) and day 12 (2.2-fold change), contrasting downregulation at other time points over the 28-day period compared with NL (Fig. 6A). Additionally, the *XTH* gene displayed upregulation under LL, with fold changes of 3.1 and 3.0 at day 20 and day 24, respectively, in comparison to other time points and NL (Fig. 6B). Finally, *SAUR19* showed upregulation under LL, with respective fold change expression values of 1.8, 3.4, and 3.7 at day 20, day 24, and day 28, distinguishing it from NL across all time points (Fig. 6C).

**Fig. 6.**
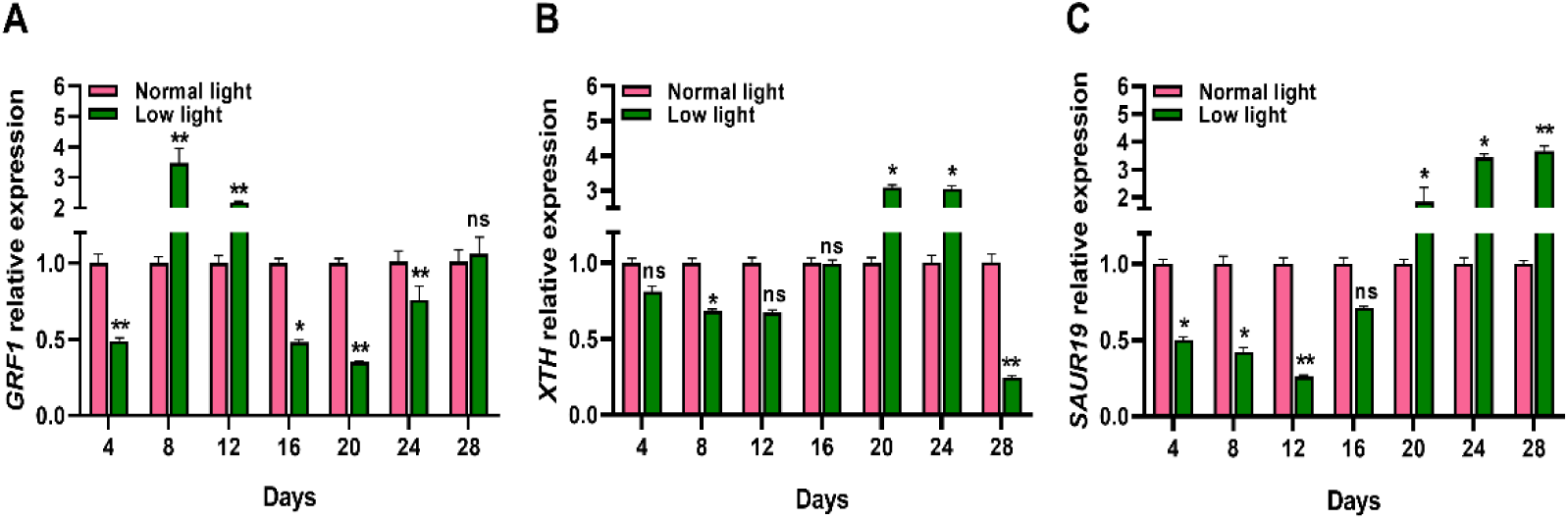
Effect of low light intensity on cell proliferation and cell expansion related gene expression. Low light make differential changes in the gene expression levels of *GRF1* (**A**), *XTH* (**B**), and *SAUR19* (**C**) over the 28-day observation period. The data is presented as mean ± SE at each time point, with * and ** indicating significant differences at p < 0.05 and p < 0.01, respectively.

## Discussion

### Leaf growth and morphology are affected by low light

Light plays a vital role in shaping the morphology and anatomical structure of plant leaves (Jiang et al. 2011). Leaf growth and development encompass a complex process characterized by well-defined cellular arrangements in lateral, longitudinal, and dorsoventral directions. Fully developed leaves exhibit three distinct asymmetrical axes: the basal-apical axis (from the leaf’s base to its tip), the ventral-dorsal axis (also known as adaxial-abaxial, where one side faces the stem as the adaxial surface and the opposite side as the abaxial surface), and the mid-lateral axis (from the leaf’s central vein to its edge) (Satterlee and Scanlon 2019). The results demonstrated that leaf area increased under low light, aligning with the findings of Yao et al (2017). This indicates that leaves possess a degree of adaptability to optimize growth in response to changing environmental conditions. Furthermore, it has been demonstrated that the leaf length-width ratio initially increased, then decreased, and eventually stabilized (Fig. S1). This indicates that during the early stages of leaf growth, transverse expansion exceeded longitudinal expansion, but as leaf development progressed, longitudinal and transverse expansion rates became roughly equal. This pattern aligns with the research of Kalve et al (2014), suggesting that later leaf growth is characterized by uniform diameter expansion, explaining the eventual elongated oval shape of the leaves. Although there were significant differences in leaf expansion rates along the lateral and longitudinal axes at the early stages of leaf development under different light intensities, leaf length showed only slight variation. Compared to normal light intensity, longer leaf expansion duration (from day 9 to day 26) was noted under low light intensity (Fig. 1), possibly related to delayed leaf senescence (low light causes premature leaf senescence) (Gad et al. 2021).

### Low light induces changes in leaf anatomy

Alterations in light intensity can influence the anatomical structure of leaves (Theroux-Rancourt et al. 2023). Lower light intensity leads to leaf thinning by reducing both the number of cell layers and cell size (Hoshino et al. 2019). This change varies across different levels of leaf organization. Results indicate that under low light, the primary difference between the dorsal and ventral axes of leaves primarily starts from palisade and spongy tissues, aligning with Fan et al. (2019) findings. This difference becomes prominent between 9 and 16 days when most mesophyll cells expand uniformly, and the intercellular space increases. Prior to this stage, (early days, from day 1 to day 9) (Fig. 3 and Fig. 4), mesophyll cells might primarily engage in cell division, with no significant differences among cell layers (Hoshino et al. 2019). Furthermore, while differences in cell division manifest during the early stages of leaf growth, variations in leaf thickness occur during the cell elongation phase, suggesting that cell elongation may play a more crucial role in the leaf thickness (Kalve et al. 2014).

### Phytohormones regulate leaf thickness

Early leaf development is an orchestrated process where plants adapt to varying light intensities by modulating their hormonal profiles. In the realm of early leaf development under low light-intensity conditions, the intricate balance of phytohormones becomes even more crucial (Ghorbel et al. 2023). This study found that IAA and GA levels were consistently higher during leaf development under low light (LL), particularly from days 4 to 20, while ABA levels were lower (days 24-28) compared to normal light (NL) conditions (Fig. 7). Under low light intensity conditions, plants prioritize a strategy characterized by higher levels of IAA and GA while maintaining lower levels of CK and ABA to optimize leaf growth and ensure efficient photosynthesis (Fan et al. 2022). Elevated IAA levels promote cell division and initiate leaf primordia (Sakamoto et al. 2022), while GA facilitates cell elongation (Tian et al. 2022), enabling leaves to grow larger, reach their proper size and shape, and capture limited available light efficiently. Conversely, CK contents were present in lower quantities during leaf development under low light (Fig. 7D), this depletion of CK might initiate a decline in photosynthetic capacity, coupled with a temporary suspension of leaf development. This mechanism ensures the redirection of energy resources towards extension growth in shaded conditions (Yang and Li 2017). In essence, these hormonal orchestrated adjustments enable plants to adapt to low light conditions efficiently. Our study confirmed the involvement of auxin, cytokinin, and gibberellin in regulating leaf anticlinal growth in cigar tobacco under low light intensity. However, it remains unclear which hormone plays the most crucial role and whether interactions among auxin, cytokinin, and gibberellin control leaf expansion in these conditions. Therefore, our next step involves investigating the effects of auxin, cytokinin, gibberellin, and their interactions on leaf expansion under low light intensities.

### Cell expansion-related gene expression is involved in the regulation of leaf thickness

The intricate regulation of cell expansion involves plant hormones, growth-regulating factors (GRFs), and other regulatory elements (Zhang et al. 2021). *GRF1*, a key player in leaf cell expansion, is evident from the contrasting sizes of leaves in the overexpression and mutant lines compared to wild types (Kim et al. 2003). Xyloglucan endotransglycosylases/hydrolases (XTHs) play a role in enhancing cell wall extensibility through the modification of cellulose and xyloglucan crosslinks. The repression of *XTH* gene expression in siz1 mutant lines, displaying defects in cell division and expansion, highlights its role in leaf cell elongation (Miura et al. 2010). SAURs, as auxin-responsive genes, act as positive effectors of cell expansion, with *SAUR19-24* specifically facilitating this process (Spartz et al. 2012). In this study, an increase was noted in UE (between day 16 and day 28—Fig. 4A), PCL (from day 10 to day 14—Fig. 4C), LE (gradual increase between day 12 to day 28—Fig. 4G), and LT (between day 10 and day 28—Fig. 4I and Fig. 3) under LL, although lower than NL. This rise could be attributed to higher expression of genes associated with cell expansion, including *GRF1* (at day 8 and day 12—Fig. 6A), *XTH* (day 20 and day 24— Fig. 6B), and *SAUR19* (from day 20 to day 28—Fig. 6C).

## Conclusions

The culmination of our research brings forth a comprehensive understanding of significant transformations unfolded in the physiology of cigar tobacco leaves under low light conditions. The interplay of light conditions intricately regulated various aspects of leaf development. Notable changes were observed in leaf morphology, anatomy, phytohormone contents, and gene expression related to cell expansion. Low light-induced alterations in leaf area, length, and width, accompanied by substantial variations in expansion rates. Anatomical analysis revealed distinct patterns in cell layers and reduced leaf thickness under low light, providing insights into the intricacies of leaf structure. Phytohormone levels, including GA, IAA, ABA, and CK, exhibited pronounced fluctuations, indicating the intricate regulatory role of light in hormonal dynamics. Gene expression related to cell expansion, such as *GRF1*, *XTH*, and *SAUR19*, demonstrated significant responsiveness to low-light conditions. This holistic understanding sheds light on the nuanced ways in which light intensity orchestrates the multifaceted processes governing leaf development in cigar tobacco plants.

## Supporting information

Fig. S1 Impact of different light conditions (intensities) on the ratio of leaf length to leaf width (LL/LW).

## Data availability

All of the data supporting the findings of this study are included in this article.

## Funding

This work was supported financially by the Agricultural Science and Technology Innovation Project of Chinese Academy of Agricultural Sciences (ASTIP-TRIC03) and the Science and Technology Program of CNTC (110202101013(XJ-05) and 110202201038(XJ-09)).

## Author Contributions

M Xinghua, R Khan, and H Gao have supervised and conceptualized the experiments related to the effect of normal light and low light on cigar tobacco morphology and anticlinal growth. X Ren and C Keling performed the experiments and compiled the data. M Xinghua, J Yang, and R Khan wrote the paper. C Yang and H Gao revised the manuscript. All authors have read and approved the manuscript.

## Competing Interests

The authors have no relevant financial or non-financial interests to disclose.

## Notes

### Competing Interest Statement

The authors have declared no competing interest.

