## Supplementary material for "Exploration of the intricacies of low light-induced changes in cigar tobacco leaf anticlinal growth: A holistic approach from anatomical and hormonal levels to gene expression": Fig. S1 Impact of different light conditions (intensities) on the ratio of leaf length to leaf width (LL/LW).

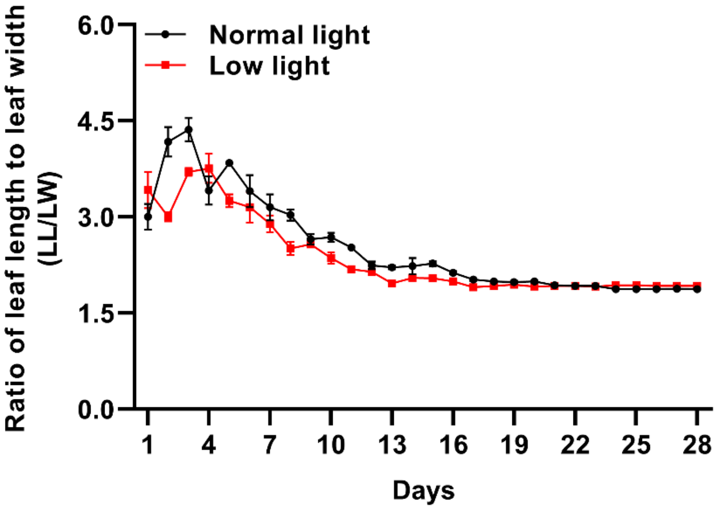


**Fig. S1** Impact of different light conditions (intensities) on the ratio of leaf length to leaf width (LL/LW). This figure represents the variation in LL/LW under normal light (300 µmol $m^{-2}$ $s^{-1}$) and low light (100 µmol $m^{-2}$ $s^{-1}$) conditions over a 28-days observation period.
